# Hourglass, a tool to mine bioimaging data, uncovers sex-disparities in the IL-6-associated T cell response in pancreatic tumors

**DOI:** 10.1101/2022.09.12.507618

**Authors:** Kazeera Aliar, Henry R. Waterhouse, Foram Vyas, Niklas Krebs, Emily Poulton, Bowen Zhang, Nathan Chan, Peter Bronsert, Sandra E. Fischer, Steven Gallinger, Barbara T. Grünwald, Rama Khokha

## Abstract

Recent advances in digital pathology have led to an explosion in high-content multidimensional imaging approaches. Yet, our ability to gainfully process, visualize, integrate and mine the resulting mass of bioimaging data remains a challenge. We have developed Hourglass, an open access user-friendly software that streamlines complex biology-driven post-processing and visualization of multiparametric data. Directed at datasets derived from tissue microarrays or imaging methods that analyze multiple regions of interest per patient specimen, Hourglass systematically organizes observations across spatial and global levels as well as within patient subgroups. Application of Hourglass to our large and complex pancreatic cancer bioimaging dataset (540,617 datapoints derived from 26 bioimaging analyses applied to 596 specimens from 165 patients) consolidated a breadth of known IL-6 functions in a well-annotated human pancreatic cancer cohort and uncovered new unprecedented insights into a sex-linked Interleukin-6 (IL-6) association with immune phenotypes. Specifically, regional effects of IL-6 on the intratumoral T cell response were restricted to male patients only. In conclusion, Hourglass facilitates multi-layered knowledge extraction from complex multiparametric bioimaging datasets and provides tailored analytical means to productively harness heterogeneity at the sample and patient level.

## Introduction

The digital pathology space is rapidly expanding and producing an immense amount of data with the potential for insights in the magnitude of OMICs profiling approaches (1). Typical sources of complex bioimaging datasets include imaging mass cytometry (IMC) and multiplexed immunohistochemistry (IHC) staining, often applied to large numbers of samples and/or tissue microarrays (TMA). In bioimaging data, one typically encounters multiple quantitative parameters per molecular target (*e.g*. percentage of positive pixels, number of positively stained cells, H scores, etc. for each stain; Table 1), as opposed to a single value per target such as obtained from OMICs experiments (*e.g*. a single protein or mRNA abundance value). The inherent multidimensionality of bioimaging data thereby poses significant challenges for targeted analysis and visualization. As a result, manual iterative hypothesis generation becomes a lengthy, labor-intensive task and a major challenge. Moreover, the ability to robustly integrate multiple parameters is equally challenging without computational knowledge yet remains indispensable for exposing meaningful biological patterns in such complex, high-dimensional data. Thus, the wealth of information that lies within these datasets is often barely utilized. In addition, bioimaging datasets and the corresponding analyses often stem from contexts with significant heterogeneity at the specimen and clinical patient levels, posing further challenges to their analysis.

Heterogeneity is a fundamental property of all tissues and oftentimes becomes exacerbated in disease conditions (2–5). Intratumoral heterogeneity is increasingly recognized as a major cancer driver (6–9), with key implications for treatment design and patient selection and a rapidly increasing appreciation of the critical disease contributions stemming from regional cellular communities (10,11). Imaging-based approaches, and thus the digital pathology field, are uniquely positioned to resolve such spatial differences down to a cellular level, enabling a more comprehensive biological understanding of the highly heterogeneous tumor ecosystem (1,12). Yet, even though standard TMA designs and typical IMC experiments incorporate multiple regions of interest in order to minimize the impact of sampling bias, the inherent opportunity of such experimental design to systematically compare and dissect regional from global biological patterns by analyzing multiple regions of the same tissue is rarely harnessed.

In order to support effective analysis of such multi-patient multi-sample bioimaging datasets, we developed Hourglass. Built as a computational toolkit, Hourglass was designed to facilitate streamlined and reproducible processing, integration, visualization and mining of multiparametric datasets, and specifically tailored to tissue-based bioimaging data. The software is available as a stand-alone R package with an optional user-friendly desktop application (at https://kazeera.github.io/Hourglass). To assess the utility of Hourglass, we queried the effects of Interleukin-6 (IL-6) in pancreatic ductal adenocarcinoma (PDAC) using an extensive dataset comprising 34 read-outs from 26 bioimaging analyses, each performed on 596 samples from 165 PDAC patients. IL-6 is a powerful regulator of tumor biology, with complex functions in both the tumor and tumor microenvironment (TME) (13,14). Several clinical trials on targeting IL-6 along with chemotherapy in pancreatic cancer are currently ongoing (e.g., NCT02767557, NCT04191421, NCT00841191). Moreover, a recent study exposed concrete effects of IL-6 on functional tumor-associated fibroblast diversity in PDAC, which has major and topical immunotherapeutic implications (15). In addition to the versatile immunomodulatory and proinflammatory effects of IL-6 in pancreatic cancer are well-established (16–18), altogether forming a strong rationale for the pursuit of combination immunotherapy/IL-6 targeting strategies in PDAC and other cancers (19–23). Notably, IL-6 levels are known to be regionally heterogeneous (19,24) with varying clinical associations, which could hint at hidden regional biological patterns evolving around this influential cytokine and this heterogeneity has potential implications for the selection of patients for future IL-6-based treatments.

We here applied Hourglass for the systematic dissection of IL-6-associated effects across multiple cellular compartments and biological processes in a well-annotated human PDAC cohort, under consideration of regional heterogeneity in IL-6 expression and clinically relevant patient subsets (sex, molecular subtypes). Our findings not only consolidate known functions of IL-6 in regulating both tumoral and stromal biology in the context human PDAC but identify a substantial involvement of regional IL-6 expression levels in distinct spatial immune phenotypes. Moreover, we discovered that the regional association of IL-6 levels with T cell communities and FAP+ fibroblasts was nearly exclusive to the male subset of PDAC patients.

## Results

### Hourglass workflow for systematic mining of annotated multiparametric bioimaging datasets

The Hourglass workflow follows three steps: dataset aggregation and import (Fig. 1A); data processing (defining comparisons, data subsets and analysis settings, selecting statistics; Fig. 1B); output mining and quality control based on visualized analysis results (Fig. 1C). For this study, we have chosen a PDAC cohort (n = 165) with detailed clinical and molecular annotations (Fig. 1A, *left*), such as transcriptional subtype, patient sex, overall survival time, tumor differentiation, etc. This cohort was used to construct a TMA, from which we generated a typical bioimaging dataset (Fig. 1A, *right*) consisting of 34 readouts from 26 stains that covered a variety of IHC markers for malignant behavior (e.g., proliferation, EMT) and TME composition (e.g., different immune cell subpopulations, fibroblasts) along with second harmonic generation microscopy and histochemical collagen stains. All stains were digitally quantified via either or both pixel- and cell-based detection metrics (Fig. S1). Where appropriate, CK19-based estimation of TME and tumor area aided compartment-specific normalization or, alternatively, quantification was restricted to manual annotations of tumoral or stromal compartments by a pathologist. Aggregation of this bioimaging data with the corresponding cohort annotations led to a final input dataset comprising 540,617 unique entries.

**Figure 1.**
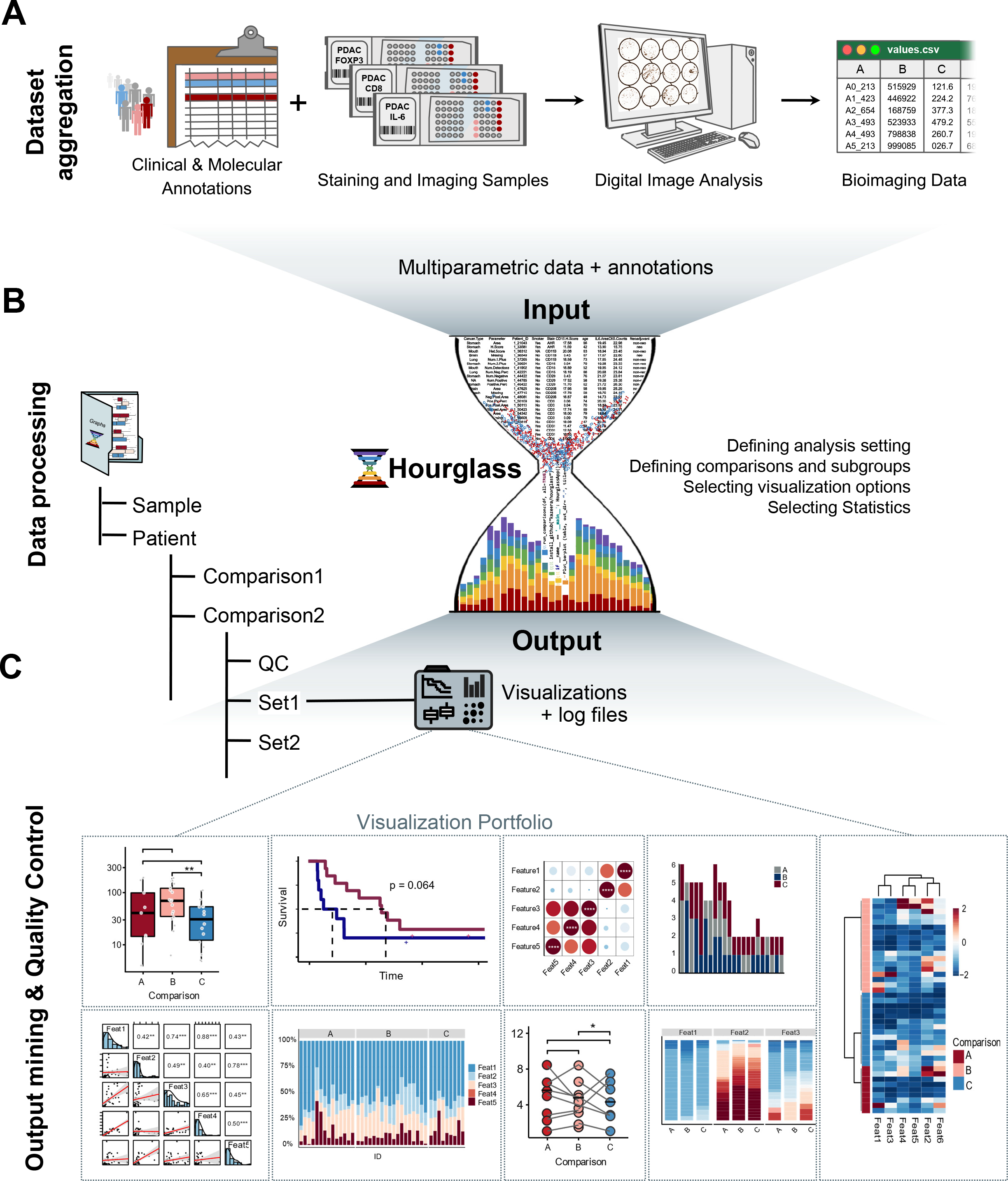
Hourglass workflow for systematic mining of annotated multiparametric bioimaging datasets. Schematic: **(A)** Multiparameteric dataset acquisition from digital bioimaging of TMA from well-annotated patient cohort, **(B)** data processing via Hourglass, and **(C)** user-selected output.

To enable the systematic and reproducible analysis of such heterogeneous multiparametric datasets, Hourglass facilitates the import of large numeric data files with extensive annotation, transforms and subsets the data based on user specifications, and produces publication-ready visualizations including pre-selected statistical tests (Fig. 1B). A variety of analysis and corresponding plot types is supported and can be selected for each analysis run; this includes comparisons between discrete groups (e.g., box plots, bar plots, slopegraphs), correlation analyses (e.g., correlation matrices, scatter plots) and Kaplan Meier survival analysis (see Table 2). The output (Fig. 1C) is stored in a structured file hierarchy with user-defined biological comparison folders and additional subfolders containing user-defined feature subsets. To enables visual data quality control and for identification of potential hidden confounders, Hourglass furthermore routinely plots all available parameters for each feature (in this context: stain) as a function of the user-defined comparison. Finally, analysis settings are logged and stored in a separate file for each run to enable full cross-run reproducibility.

### Graphical user interface (GUI) supports navigation through Hourglass workflow without prior computational expertise

The software application for Hourglass was made accessible to non-expert users, while also supporting customizable analysis at the experienced user level. Therefore, a graphical user interface (GUI) was built to take users through the Hourglass algorithm via step-by-step tabs (Fig. 2A-C). In brief, the Hourglass application allows manual selection of input data and defining biological comparisons of interest based on values present in the input data files (Fig. 2A), selection of feature subsets and customization of colors (Fig. 2B), along with advanced options to iteratively refine analyses and a final selection of parameters for the current run, including manual selection from supported plot types and quality control settings (Fig. 2C). The application then saves the user analysis specifications (Fig. 2D) as an Excel file and interfaces to the R statistical environment. Finally, the corresponding Hourglass R package produces the output as a sequential background process (Fig. 2E). Computational expert users can work directly within the Hourglass R package, without the GUI.

**Figure 2.**
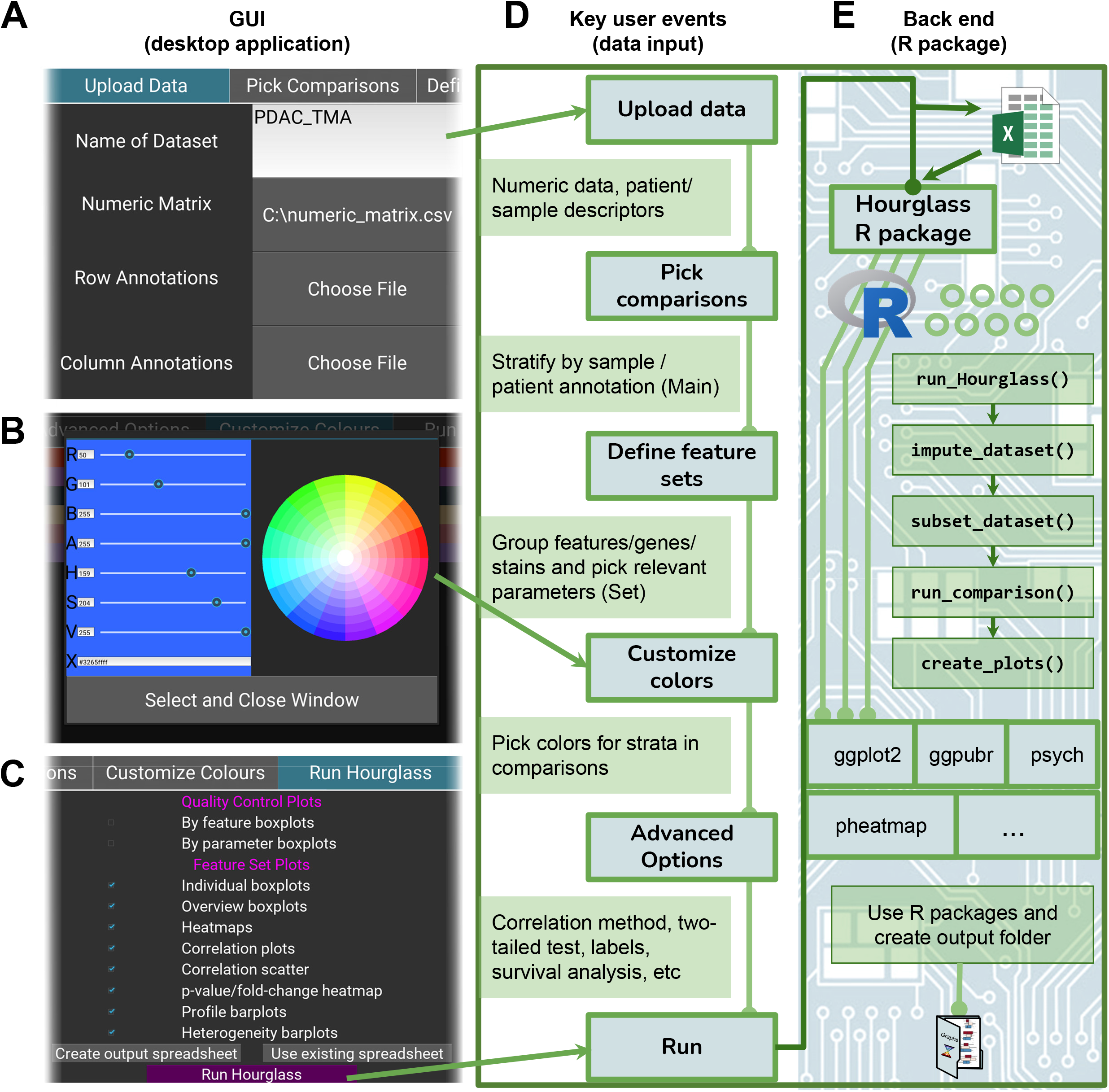
Graphical user interface supports navigation through Hourglass workflow. GUI tab layout supporting **(A)** data import **(B)** customization of colors for graphs, and **(C)** user-selected options. **(D)** Key steps in analysis pipeline corresponding to tabs in GUI. **(E)** Run button logs user-defined data file paths and choices to Excel file and submits to R package to create output folder via R functions and dependencies.

### Inter- and intra- tumoral heterogeneity in IL-6 expression levels

To test the utility of Hourglass, we chose to profile the breadth of IL-6-related biology in human PDAC, aiming at a systematic dissection of the powerful multipronged effects under consideration of the known intratumoral heterogeneity of IL-6. An IL-6 IHC stain was optimized using a set of control tissues (Fig. S2A) and then applied to a TMA from well-annotated PDAC patients (Fig. 1A). IHC showed varying IL-6 expression levels across PDAC patients with staining in malignant epithelial cells, fibroblasts and immune cells (Fig. 3A, *left*). Furthermore, IL-6 levels varied intratumorally within both malignant epithelial and stromal regions (Fig. 3A, *right*). As a result, when we divided all samples based on 33% quantiles (IL-6^lo^, IL-6^int^ and IL-6^hi^ Fig. S2B), we observed that different TMA cores from the same patient exhibited varying IL-6 expression categories (Fig. 3B) in 33.6% of PDAC cases. While our analysis here focuses on IL-6, we observed similar levels intratumoral heterogeneity (i.e., multiple cores from the same tumor being classified as IL-6^lo^, IL-6^int^ and/or IL-6^hi^) for a range of markers in our staining panel (e.g., CD3: 46.5%; CD68: 42.2%; αSMA: 52.2% of cases). Altogether, IL-6 was commonly expressed in human PDAC and exhibited a spatially heterogeneous expression pattern.

**Figure 3.**
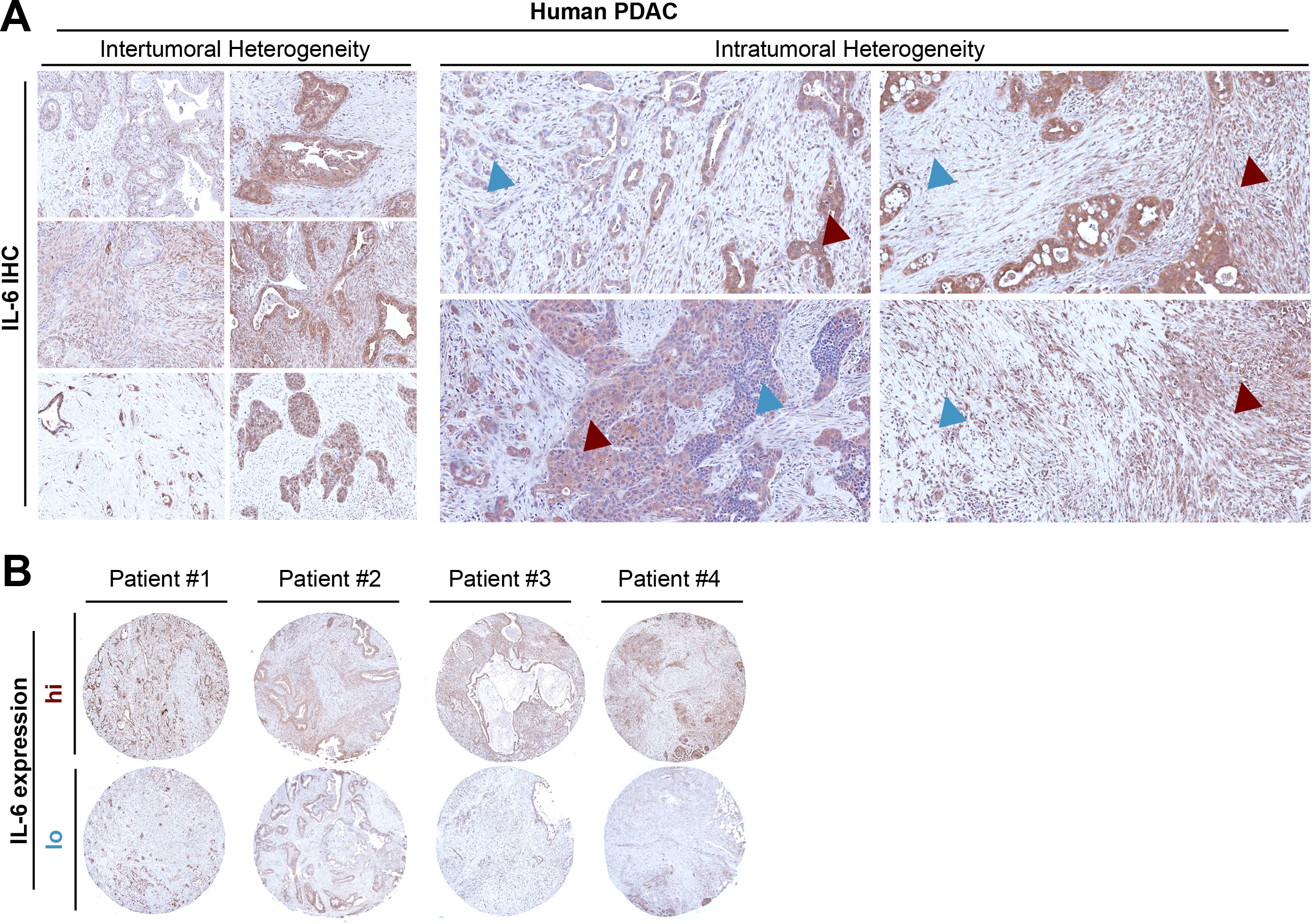
Inter- and intra-tumoral heterogeneity of IL-6 expression levels in PDAC. **(A)** Representative images of human PDAC tumors showing varying intertumoral (left) and intratumoral (right) IL-6 levels; arrowheads indicate IL-6 low (*blue*) vs. high (*red*) regions. **(B)** Varying IL-6 expression within different TMA cores from four representative PDAC tumors.

### IL-6^high^ PDAC patients show distinct myeloid, endothelial and ECM properties but little involvement of lymphoid cells

Bioimaging data from TMAs or other multiple regions of interest (ROI)-based methods are typically collapsed to the patient level, i.e., expression levels from multiple sampled regions are averaged to achieve robust patient stratification (Fig. 4A). This common analysis standard was implemented in Hourglass and served as entry point for investigation of the functions of IL-6 in PDAC. First, we compared PDAC patients by IL-6 expression quantiles (Fig. S3A) across our full stain panel (Fig. 4B) to get an overview of the involvement of IL-6 in human PDAC disease biology.

**Figure 4.**
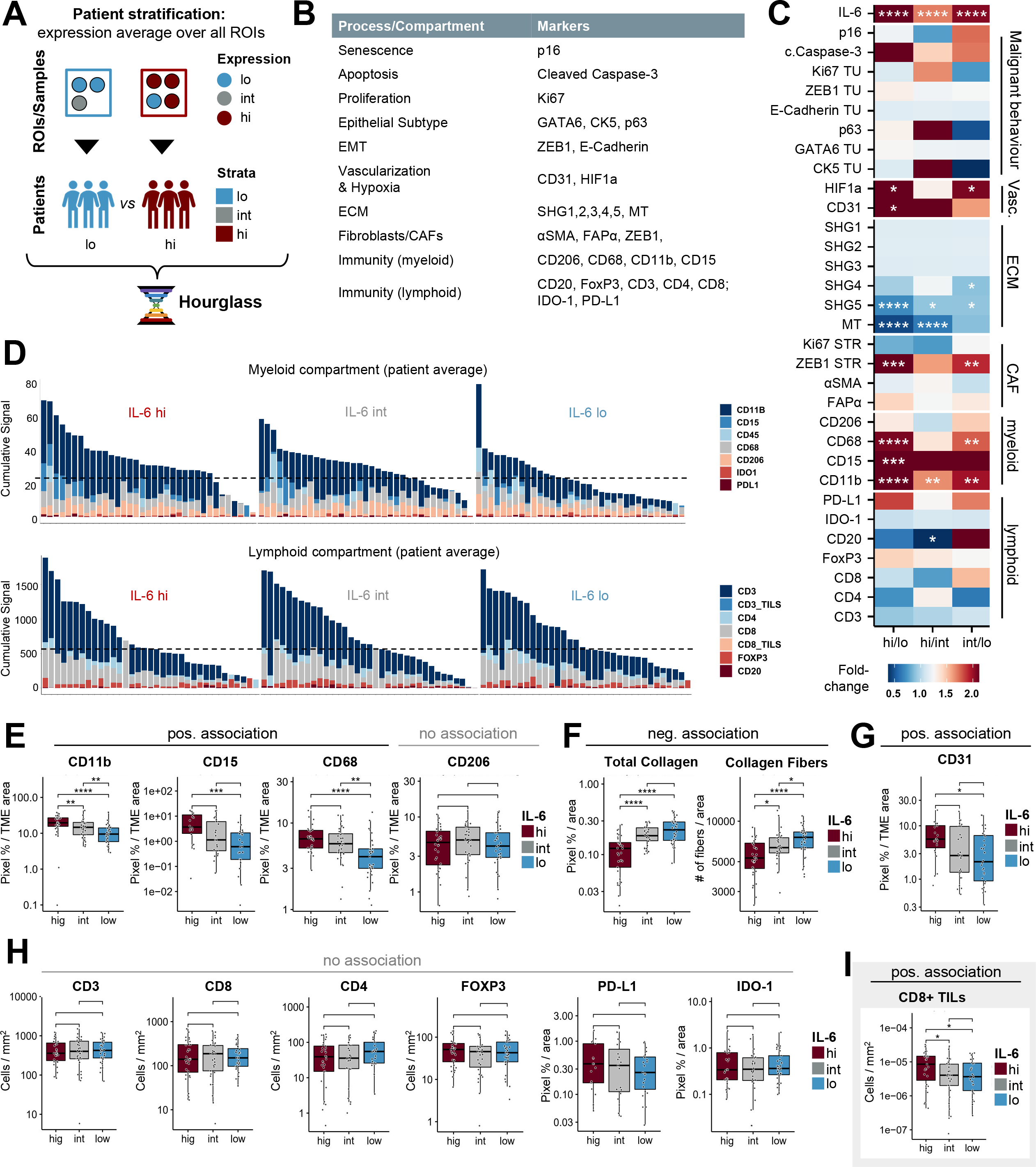
IL-6^hi^ PDAC patients show distinct myeloid, endothelial and ECM properties but little involvement of lymphoid cells. **(A)** Schematic: Hourglass can average and compare IL-6 expression over multiple samples/regions of interest (ROIs) per patient. **(B)** Staining panel used in this study. **(C)** Heatmap of fold-change expression and significance levels for all stains in (**B**) across IL-6^hi^ (hi), IL-6^int^ (int) and IL-6^lo^ (lo) expressing human PDAC tumors. SHG1: Mean Col Width, SHG2: Mean Col Straightness, SHG3: Mean Col Length, SHG4: Col Alignment, SHG5: Number Col (where Col is Collagen Fiber). **(D)** Composite plot of all myeloid (*top*) and lymphoid (*bottom*) marker quantification results per patient grouped by IL-6 levels (hi/left, int/middle, lo/right). **(E)** Individual digital quantification results of IHC stains for CD11b, CD15, CD68, CD206 (myeloid markers), **(F)** SHG1, MT (ECM), **(G)** CD31 (vascularization), **(H)** CD3, CD8, CD4, FOXP3, PD-L1 (lymphoid), as well as manual counts of **(I)** tumor infiltrating CD8+ lymphocytes (CD8+ TILs). **(F)** Number of collagen fibers as assessed through second harmonic generation microscopy. **(E-I)** Boxplots compare IL-6^high^ (hig), IL-6^int^ (int) and IL-6^low^ (low) patients. **(C, E-I)** n = 143; Wilcoxon test. *p < 0.05, **p < 0.01, ***p < 0.001,****p < 0.0001.

At the patient level (Fig. 4C), IL-6 showed no significant associations with malignant cell behavior yet was strongly linked to changes in the TME. Specifically, high IL-6 levels were associated with increased vascularization, lower ECM content, and increased presence of the neutrophil marker CD15, monocyte marker CD11b and macrophage marker CD68 but not the M2 phenotype marker CD206. In contrast, CAF markers such as αSMA and FAPα were not significantly altered and neither were T cell markers (CD3, CD4, CD8, FoxP3) nor checkpoint molecules (PDL1, IDO1). Accordingly, composite analysis of all myeloid and all lymphoid markers confirmed an overall enrichment of myeloid but not lymphoid cells in human PDAC tumors along with increasing IL-6 expression (Fig. 4D). Inspection of the individual datapoints further showed a clean dose-dependent associations with IL-6 for the following markers: CD15, CD11b and CD68 (higher along with IL-6; Fig. 4E), total collagen content and number of collagen fibers (lower with high IL-6; Fig. 4F), and vascularization (CD31, HIF1; higher along with IL-6; Fig. 4G). The B cell marker CD20 showed a weak, non-dose-dependent association as it was lower in IL-6^hi^ compared to IL-6^Int^ patients (Fig. S3B). Altogether, IL-6^hi^ PDAC tumors exhibited increased myeloid cell population and vascularization along with reduced ECM content.

Intriguingly, none of the T cell marker or checkpoint molecule stains showed any association with IL-6 levels and none except PDL1 even trended along with IL-6^hi^ levels (Fig. 4H). This was unexpected given the well-established and central role of IL-6 in T cell functionality, expansion and survival in a variety of pathological contexts and its shown benefits in combination with immunotherapy (16,19,25–28). We thus performed a targeted manual analysis to count tumor infiltrating CD8+ lymphocytes (CD8 TILs) in direct physical contact with malignant epithelial cells, which is indicative of the presence of functional CD8 effector T cells. This indeed revealed a small but significant increase in CD8+ TILs in IL-6^hi^ PDAC patients (Fig. 4I). In light of the known regional heterogeneity in the intratumoral T cell response in PDAC (10) we decided to explore this observation further in the regional analysis formats that are supported by Hourglass.

### Intratumoral heterogeneity-informed Hourglass workflow links IL-6 expression to regional T cell enrichment

To complement the patient-level analysis (Fig. 4A), Hourglass was designed to support an otherwise identical analysis in parallel at the individual sample level (*i.e*., by-sample (Fig. 5A). Notably, in this regional analysis, IL-6^hi^ samples showed a tendency to spontaneously group together across all stains (Fig. 5B), suggesting IL-6 is overall involved in tissue composition. As before at the patient level (Fig. 4C), high regional IL-6 was strongly linked to differences in TME composition, associating with increased myeloid cells and endothelium, and decreased ECM content (Fig. 5C). Only the B cell marker CD20 did not associate with regional IL-6 levels (Fig. S4). Beyond these differences, this analysis level revealed higher numbers of FAP+ CAFs, as well as an increase in CD3+, CD8+ cells and checkpoint molecules IDO-1 and PD-L1 in IL-6^hi^ regions in human PDAC. Accordingly, composite analysis of the myeloid and lymphoid cells showed an overall enrichment of both immune compartments in regions with high IL-6 expression in human PDAC (Fig. 5D), which had not been apparent in the global analysis (Fig. 4D). Inspection of the individual datapoints also showed the same clean expression-dependent associations with myeloid markers (CD11b, CD15, CD68; Fig. 5E), ECM content (Fig. 5F) and vascularization (CD31; Fig. 5G) as before. In addition, there was a clear association of regional IL-6 levels with CD3+ and CD8+ T cells as well as PDL-1 and IDO1 (Fig. 5H) and a prominent increase in CD8 TIL counts (Fig. 5I).

**Figure 5.**
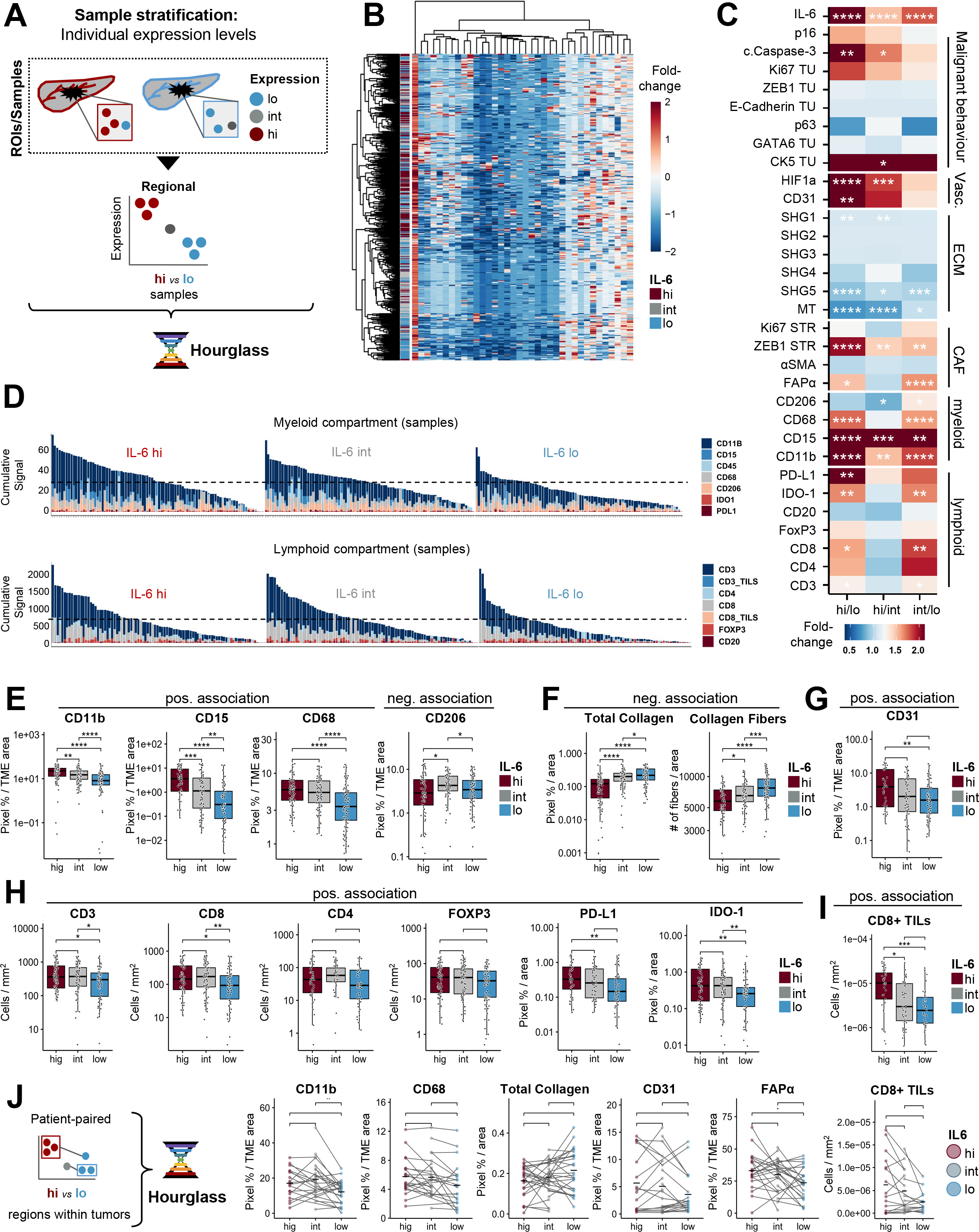
Intratumoral heterogeneity-informed Hourglass workflow links IL-6 expression to regional T cell enrichment. **(A)** Schematic: Hourglass can compare IL-6 expression in individual samples/ROIs. **(B)** Heatmap visualizing hierarchal clustering of markers in staining panel in all samples (n= 516), grouped by IL-6 level. **(C)** Heatmap of fold-change expression and significance levels for all stains across IL-6^hi^ (hi), IL-6^int^ (int) and IL-6^lo^ (lo) samples. SHG1: Mean Col Width, SHG2: Mean Col Straight, SHG3: Mean Col Length, SHG4: Col Alignment, SHG5: Number Col (where Col is Collagen Fiber). **(D)** Composite plot of all myeloid (*top*) and lymphoid (*bottom*) marker quantification results per sample grouped by IL-6 levels (hi/left, int/middle, lo/right). **(E)** Individual digital quantification results of IHC stains for CD11b, CD15, CD68, CD206 (myeloid markers), **(F)** SHG1, MT (ECM), **(G)** CD31 (vascularization), **(H)** CD3, CD8, CD4, FOXP3, PD-L1 (lymphoid), as well as manual counts of **(I)** tumor infiltrating CD8+ lymphocytes (CD8+ TILs). **(F)** Number of collagen fibers as assessed through second harmonic generation microscopy. **(E-I)** Boxplots compare IL-6^high^ (hig), IL-6^int^ (int) and IL-6^low^ (low) samples. **(J)** Schematic: Hourglass can compare difference IL-6 levels within the same patient (left); Patient-paired slope-graphs show CD11b, CD68 (myeloid), FAPα (CAF), MT (ECM) CD31 (vascularization), and CD8+ TILS levels in IL-6^high^+ IL-6^int^ (hig) and IL-6^low^ (low) samples within the same patient (right). **(C, E-J)** n = 516.Wilcoxon test. *p < 0.05, **p < 0.01, ***p < 0.001,****p < 0.0001.

To test if these additional and/or stronger trends were merely a reflection of increased sample numbers, we compared paired IL-6^hi^ vs IL-6^lo^ regions within the same tumors, leveraging the presence of IL-6^hi^ and IL-6^lo^ cores from the same patients in our TMA (Fig. 3B, Fig. 5J *left*). This showed that regional differences in intratumoral IL-6 expression trend to be directly accompanied by regional enrichments in myeloid cells, endothelial cells, FAPα+ (but not αSMA+) CAFs, tumor-infiltrating CD8+ T cells, and a decrease in ECM (Fig. 5J). Altogether, regional variation in IL-6 expression appears to have distinct consequences for local TME composition, which can be masked when averaging regions of various expression within patients.

### Systematic comparison of IL-6 effects between patient subsets identifies sex-specific immune phenotypes in PDAC

PDAC is clinically heterogeneous and biological patterns might vary between patient subgroups. Iterative refinement of analyses to identify confounder subsets can thus be instrumental to understand underlying biology. For instance, molecular subtypes of PDAC (29) differ in their baseline immune microenvironments (30,31) while IL-6 itself has been implicated in sex-differences in the context of hepatocellular carcinoma (32). To test whether the involvement of IL-6 in key biological patterns (immune phenotypes, TME composition) was confounded within specific PDAC patient subsets, we applied parallel subgroup comparison under otherwise identical analytical conditions across global and regional analysis levels, leveraging all steps in the Hourglass workflow (Fig. 6A).

**Figure 6.**
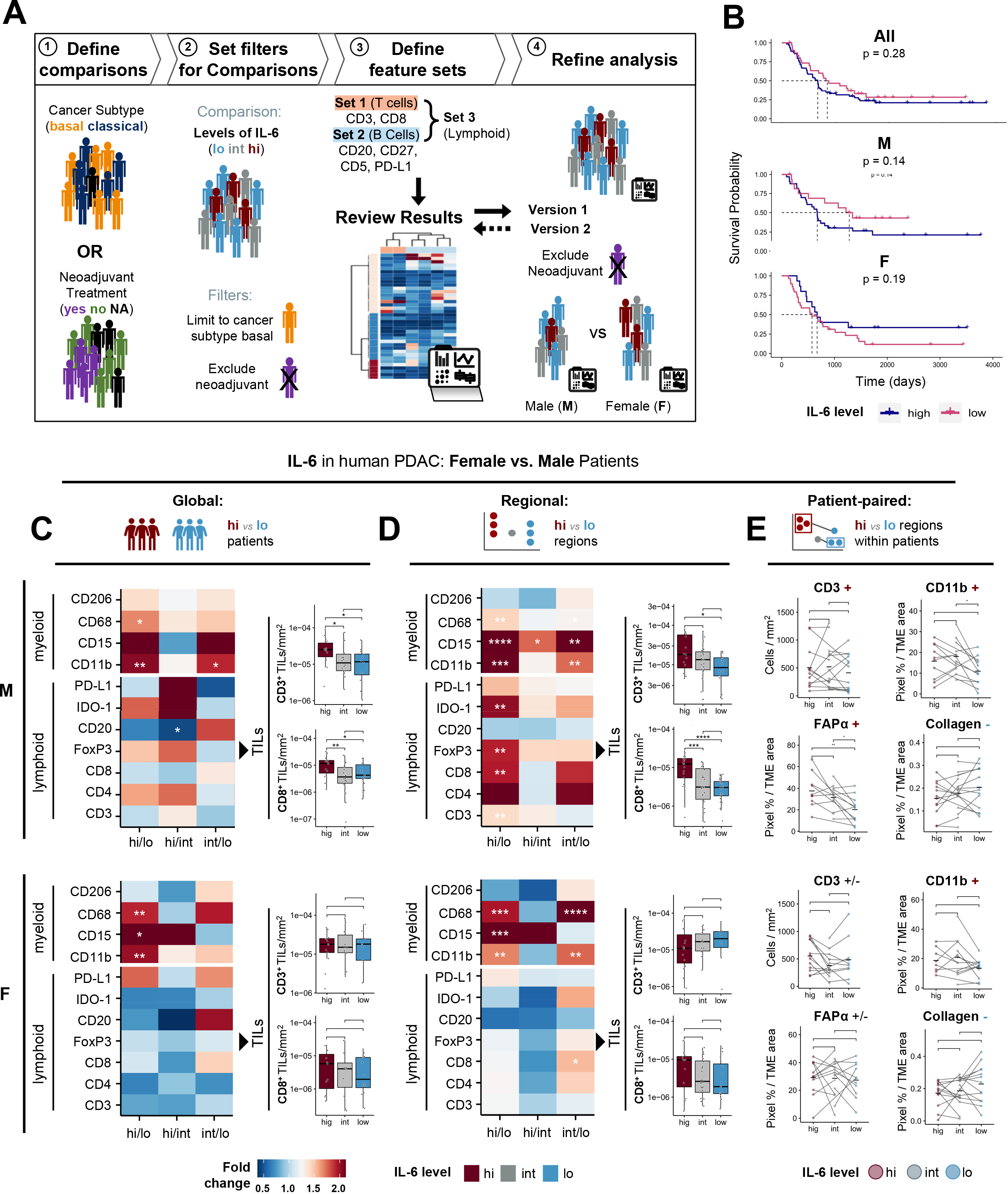
Systematic comparison of IL-6 effects between patient subsets identifies sex-specific immune phenotypes in PDAC. **(A)** Schematic: Hourglass systematically dissects observations across patient subgroups within user-defined biological sets of features/stains. **(B)** Kaplan-Meier analysis of overall survival time in all patients (left), and basal-like (middle), classical (right) sub-cohorts; strata: IL-6 level; log-rank tests. M, male; F, female; **(C-E)** Comparison of IL-6 effects in male (top panels) versus female (bottom panels) PDAC patients across global (**C**), regional (**D**) and patient-paired (**E**) levels. (**C, D**) Heatmaps depicting foldchange expression and significance levels for selected immune markers; boxplots show manual TIL counts. **(E)** Patient-paired slope-graphs depicting CD3 (T cells), CD11b, (myeloid), FAPα (CAF), MT (ECM) and CD8+ TILS levels. (**C-E**) All levels are stratified by IL-6 expression quantiles into IL-6^hi^ (hi), IL-6^int^ (int) and IL-6^lo^ (lo) groups. Wilcoxon test. *p < 0.05, **p < 0.01, ***p < 0.001,****p < 0.0001.

While IL-6 expression levels were the same in male and female patients (Fig. S5A), we noticed that male patients trended towards worse survival outcomes as a function of high IL-6 expression (Fig. 6B). At the global level (all samples averaged by patient), this was accompanied by similar myeloid cell involvement in male vs female IL-6^hi^ patients (Fig. 6C, *heatmaps*) but a significant increase in both CD3 TILs and CD8 TILs that was absent in female patients (Fig. 6C, *boxplots*), despite a trend towards higher CD8 TILs that remained non-significant. Strikingly, at the regional level (samples divided by IL-6 expression levels), we again observed a consistent increase of T cell markers (CD3, CD8, CD4, FoxP3) and checkpoint molecules (PDL1, IDO1) as well as CD3 TILs and CD8 TILs in IL-6^hi^ PDAC regions yet this was entirely restricted to male patients (Fig 5H-I vs. Fig. 6D). This indicates the presence of a male-specific regional T cell response linked to IL-6 expression levels in human PDAC. Moreover, when we compared the TME composition of paired intratumoral IL-6^hi^ and IL-6^lo^ regions between sexes (Fig. 6E), we observed larger scale differences: in male patients, IL-6^hi^ compared to IL-6^lo^ regions exhibited more CD3 T cells, more CD11b myeloid cells, more FAPα+ CAFs, and less ECM content. In female patients, on the other hand, we observed unaltered CD3 T cells, and unaltered FAP+ CAFs, while the reduced ECM content and the increase in CD11b myeloid cells were consistent between both sexes. Altogether, this suggests that regional variations in IL-6 levels within human PDAC lead to spatial effects on TME composition that are in part sex-specific, namely with male TMEs being overall more inflamed and more CAF-activated in the presence of IL-6.

### IL-6 effects vary only slightly between molecular subtypes of PDAC

We then proceeded to test whether there are differences in the IL-6 response between PDAC basal-like and classical molecular subtypes (29), since we had noticed that IL-6 expression was higher in classical than in basal-like PDAC tumors (Fig. S6B). Patients with basal-like tumors exhibited significantly worse survival outcomes as a function of high IL-6 expression (Fig. S6C). At the global level (all samples averaged by patient), this was accompanied with a much less pronounced myeloid involvement in IL-6^hi^ regions of basal-like than of classical PDAC tumors (Fig. S6D, *heatmaps*). We in turn observed higher CD3 TILs and CD8 TILs in IL-6^hi^ regions of basal PDAC tumors (Fig. S6D, *boxplots*) even though classical PDACs trended in the same direction. Yet, at the regional level (samples by IL-6 expression levels), both basal and classical tumors exhibited similar differences in response to IL-6 across lymphoid and myeloid markers as well as CD3 TILs and CD8 TILs (Fig. S6E). Likewise, when we compared the TME composition of paired IL-6^hi^ and IL-6^lo^ regions within tumors of different subtypes (Fig. S6F), we observed very similar effects across both subtypes: unaltered T cells (CD3), increase FAP+ CAFs and decreased ECM; only myeloid cells (CD11b) were more strongly associated with IL-6 levels in the classical subset of PDAC patients.

Altogether, this systematic analysis of IL-6 in human PDAC through the Hourglass computational toolkit has revealed that a) T cell responses in PDAC were linked to IL-6 expression levels but largely occur as a regional phenomenon, b) regional IL-6 dependent T cell responses in human PDAC were largely restricted to male patients and c) higher IL-6 expression was associated with worse survival outcomes in basal-like subset of PDAC patients, which was however not directly linked to major differences covered by our tumor biology staining panel.

## Discussion

Technological advances in biomedical research are producing immense amounts of data and the resultant downstream analysis can become a formidable undertaking. To address this obstacle, we constructed a framework for systematic interrogation of multiparametric datasets. The software package, called Hourglass, is informed by tissue heterogeneity and able to strategically process multiparametric datasets with a focus on functionalities that are frequently required for bioimaging datasets, such as assembly of feature subsets and regional versus global analysis levels. We showcase its utility for knowledge extraction from a PDAC TMA dataset, where it derives novel insights into IL-6-related regional biology and the identification of clinically relevant patient subsets in the context of future IL-6 directed immunotherapeutic strategies for PDAC.

Digital image analysis, as opposed to semi-quantitative scoring, is becoming a common standard and has significant benefits in terms of larger scale experiments such as a wide staining panel applied to a TMA, normalization and implementation on analysis standards across labs and institutions, and cost-effectiveness. (33–36) Hourglass was designed as a user-friendly and robust analysis tool to systematically and reproducibly mine large bioimaging datasets in the rapidly expanding digital pathology space. Naturally, alongside the boom of big data in biomedical and clinical spaces, tools are constantly created to query public and private datasets. However, there are very limited tools that facilitate complex computational bioimage analysis in a streamlined approach. This is due to several reasons, but most notably, the majority of intuitive and/or interactive tools are geared towards omics analysis and consequently provide little to no support for complex multiparametric data structures. At the same time, computational experts in the bioimaging space are mostly engaged in rather high-end artificial intelligence-based analytical approaches (36–38), which are undoubtedly extremely powerful but not well-suited to generate software solutions suitable for systematic data mining of standard bioimaging applications. Consequently, we hope that Hourglass will particularly appeal to non-expert users and encourage more iterative and heterogeneity-informed mining of large bioimaging dataset with extensive annotations, as it supports customizable, iterative, and reproducible data exploration via a streamlined and semi-automated workflow.

Careful analysis of multiparametric bioimaging datasets bears specific demands and challenges, such as a high need for quality control across sample due to sample dropouts and/or staining artifacts. In addition, with multiple options for how to analyze different stains (cell-based versus pixel-based metrics, weighted scoring such as allred or H-scores), the ability to systematically compare and benchmark multiple parameters for each stain and select the appropriate one is essential for optimal analyses. Yet, this becomes laborious and error-prone if done manually across multiple stains. Hourglass has thus been tailored to address specific needs of both multiparametric data (manual parameter selection, parallel comparison under otherwise identical conditions even within larger stain panels) and bioimaging datasets (integration and subsetting of large stain panels, heterogeneity-informed navigation of regional and global levels, quality control across all parameters per stain). In addition, Hourglass supports various required data post-processing steps that might be particularly challenging for computational non-expert users. For example, to account for dropouts (*e.g*., insufficient staining quality, cores lost at washing steps, tissue folds), Hourglass can perform imputation of missing data to then facilitate downstream visualization per unsupervised clustering in heatmaps and correlation analyses. Based on user specifications, Hourglass then aggregates and visualizes data subsets and applies statistical testing. Outputs are structured in a coherent file hierarchy and analytical specifications are logged, to facilitate iterative hypothesis generation and testing. We are to the best of our knowledge not aware of any other software applications that provide such a comprehensive toolkit tailored towards relevant functionalities for bioimaging and multiparametric datasets to date.

Tissue heterogeneity and regional cell communities are increasingly understood as a relevant disease-driving features rather than noise (10,11). Nevertheless, the most common approach to preparing bioimaging data from TMAs is to average over multiple samples or ROIs per patient while a whole wealth of information may be contained within those multiple individual regions per tumor. Hourglass was designed to access all levels of this unique data structure. Specifically, we harness this standard redundant sampling approach taken in TMAs and other multi-ROI based bioimaging approaches to systematically dissect regional from inter-patient heterogeneity and generate a new level of insight into tissue biology from the otherwise unused or discarded information. In essence, Hourglass incorporates different analytical levels in its heterogeneity-informed approach by supporting parallel analysis of patients, individual samples, and patient-paired samples compared as a function of a parameter of interest (here, IL-6 expression). We showcase how such systematic dissection of intra- and intertumoral heterogeneity can lead to new insights into disease-relevant traits and clinically relevant IL-6 biology. Notably, these different levels of analysis can be used alone or in combination, *i.e*. users can focus on the standard by-patient analysis only or perform the analysis under identical settings at the global, regional and patient-paired levels to systematically identify commonalities and differences. Altogether, we strongly recommend exploring all levels first and then decide based on the obtained results. For instance, if regional differences are driving global trends, as we have observed for IL-6 in male vs female PDAC case, then heterogeneity or sampling bias is highly relevant in this context. On the other hand, when regional analyses do not provide additional insight such as in the case IL-6 in the context of basal-like vs classical PDAC, subsetting may simply not be warranted for this specific research question.

IL-6 is a key therapeutic target, not only in PDAC but many inflammation-related diseases (39) and affects essentially all aspects of malignant biology. Yet, there are numerous conflicting reports as to whether IL-6 levels are a prognostic indicator in PDAC and other cancers, and these varying outcomes suggest significant clinical heterogeneity in the effects of IL-6. In addition, IL-6 expression is known to be regionally heterogeneous (19,24), posing challenges for sampling and due to spatial confinement of key biological associations that might be missed in current standard approaches. By systematically navigating through all these layers of complexity, Hourglass revealed novel, largely sex-specific, spatial effects of IL-6 expression on TME composition in human PDAC. In specific, we found that T cells, as opposed to the major myeloid cell populations, responded to regional variation in IL-6 expression, and these were masked when averaging multiple regions of various expression within all patients. Moreover, this regional effect of IL-6 on the T cell compartment was almost entirely restricted to male patients. Notably, we also observed that FAP+ fibroblasts, another key stromal cell population with presumed immune-modulatory functions (40) were also restricted to these IL-6^hi^ regions in male patients only. In turn, there was a consistent regional and global anticorrelation of IL-6 with the amount of ECM across sexes and subtypes. Notably, CD8 TILs were the only T cell subset that fairly consistently aligned with IL-6 levels across multiple patient subsets, hinting at a more robust link between IL-6 and CTL functionality. Altogether, this exposes novel associations of IL-6 with TME biology in human PDAC, sex-disparities within the lymphoid arm of the intratumoral immune response and provides a systematic roadmap for navigating such complex phenomena across spatial levels and patient subsets. The here-discovered sex disparities in the IL-6 related intratumoral T cell response in PDAC have furthermore major implications for the design of immunotherapeutic strategies and the selection of patients for such new treatments. Moreover, such disparities in patient subgroups might in part be responsible for the conflicting clinical correlates of IL-6.

## Supplemental Figures

**Figure S1. Quantification metrics for digital IHC images, related to Figure 1**

Where applicable, stains were digitally quantified using QuPath via either or both cell-(*left*) and pixel-(*right*) based detection methods.

**Figure S2. Intratumoral heterogeneity of IL-6 and other stains within patients, related to Figure 3**

**(A)** IL-6 staining controls (kidney, liver, pancreas); representative images.

**(B)** Binning of human PDAC samples based on tertiles of IL-6 expression.

**Figure S3. Expression of IL-6 and CD20 across patients, related to Figure 4**

**(A)** Binning of human PDAC tumors based on tertiles of IL-6 expression.

**(B)** Digitally quantified CD20 IHC stain results plotted as function of IL-6 expression per tumor.

**Figure S4. Expression of CD20 across samples, related to Figure 5**

Digitally quantified CD20 IHC stain results plotted as function of IL-6 expression per sample.

**Figure S5. IL-6 effects vary only slightly between molecular subtypes of PDAC, related to Figure 6**

Boxplots show expression of IL-6 in patients across **(A)** sexes and **(B)** Moffitt molecular subtypes.

**(C)** Kaplan-Meier analysis of overall survival time in all patients (*left*), and basal-like (*middle*), classical (*right*) sub-cohorts; strata: IL-6 level; log-rank tests.

**(D-F)** Comparison of IL-6 effects in PDAC patients with basal-like (top panels) versus classical (bottom panels) transcriptomic subtype across global (**D**), regional (**E**) and patient-paired (**F**) levels. (**D, E**) Heatmaps depicting foldchange expression and significance levels for selected immune markers; boxplots show manual TIL counts. **(F)** Patient-paired slope-graphs depicting CD3 (T cells), CD11b, (myeloid), FAPα (CAF), MT (ECM) and CD8+ TILS levels. (**D-F**) All levels are stratified by IL-6 expression quantiles into IL-6^hi^ (hi), IL-6^int^ (int) and IL-6^lo^ (lo) groups. Wilcoxon test. *p < 0.05, **p < 0.01, ***p < 0.001,****p < 0.0001.

## Methods

### Resource Availability

#### Lead contact

Further information and requests for resources and reagents should be directed to and will be fulfilled by the lead contact, Rama Khokha.

#### Materials availability

This study did not generate new materials.

### Data and code availability

Hourglass source code is available on Github, see https://kazeera.github.io/Hourglass for user manual.

### Method details

#### Acquisition of pancreatic cancer tissue microarray with immunohistochemistry

Hourglass was built around the PDAC IHC TMA dataset, presented in (10). Briefly, pancreatic cancer tissues were obtained from patients, samples were placed as cores (diameter 1.2 mm) into TMAs, 5 μm lateral sections, and stained for different genes or features. Pixel-based or cell-based detection parameters were manually set and optimized for each stain. Quantification results were quality controlled by visual inspection of the 10 cores with the highest detected staining levels and the 10 cores with the lowest detected staining levels for each slide for each stain. Next, images of slides were quantified by Numeric readouts or parameters acquired for each stain for each core, for example the area, number of positive pixels, etc. Next, a merging script was used to merge all the stain files into one table and integrated with patient/sample annotations information including age, sex, cancer subtype, treatment history, mutation statuses. Each region of interest had a unique identifier containing the core location on the TMA grid, slide, links to other dataset acquisition (e.g. sequencing) and links to clinical and histopathology. Each patient has a unique case ID and 2-8 samples/regions of interest.

#### Description of bioimaging data

The dataset used in this study consists of 596 cores or tissue samples from 165 patients (i.e. 1 to 15 samples per patient; average 3.6), first published in (10). Each sample had 26 stains applied, including histochemical stains, immunohistochemical stains, and second harmonic generation microscopy. Digital quantification of these stains using the bioimage analysis softwares QuPath and CT-FIRE recorded 22 unique parameters (Table 1).

The final *sample x feature-parameter* matrix dimensions are 648 x 696, resulting in 451,008 data points. In addition, patient data was used to define multiple recorded subgroups, such as age, cancer subtypes, and treatment history that could be compared (represented as a clipboard in Fig. 1A). In our dataset, 50 and 59 annotations were labeled for patients and for samples respectively.

#### Statistical Analyses

Where appropriate, the quantitative analyses are described in the relevant sections of the Method details. Statistical and visual analyses were performed using Hourglass and R (v4.1.0). Quantitative variables were compared by Wilcoxon rank sum test, pairwise in slopegraphs. Fold-changes were calculated by dividing means of respective groups. Survival curves were plotted by the Kaplan–Meier method and p values were assessed based on logrank test. The specific statistical tests used are indicated in the figure legends. All tests were two-sided. Statistical significance was set at p<0.05. Correlation coefficients were determined by the Spearman method.

#### Development of R package

The standalone Hourglass R package was built using R v4.1.0. We compiled a number of existing data filtering functions (such as subset retrieval, imputation and outlier removal) and plotting functions and wrote our own.

#### Development of graphical user interface

The Kivy design language (v2.0.0) and pyinstaller (v4.8) in python3 was used to create a standalone executable.

#### Details of IL-6 Hourglass run

For this study, we customized an Hourglass analysis to interrogate the role of IL-6 in PDAC with 8 main comparisons (comparing subgroups in samples and patients; e.g. looking at IL-6 in basal vs classical tumors), 22 feature sets (grouping stains; e.g. vascularization, immune, epithelial behavior) and 10 plot types. In parallel, Hourglass imputed and ran the same comparisons on this version of the dataset. Hourglass ran as a background sequential process for 7.393 hours on a Windows System (32GB RAM, Intel (Alt-0174) ^Ⓡ^ Core™(Alt-0153) i7 CPU). The result was an output folder with a total of 17,386 PDF files contained in subfolders with relevant biological analysis, amounting to 3.40 GB (3,401,516,890 bytes) of publication-ready plots.

## Acknowledgments

This work was supported by grants to RK from CCS and CIHR. BTG held fellowships from the Princess Margaret Foundation, EMBO (ALTF-116-2018), Alexander von Humboldt Foundation (DEU-1199182-FLFP), and CRS (Next Generation of Scientists Scholarship). We thank Parinaz Nasr Esfahani for supporting software development.

The authors declare no competing financial interests.

## Author contributions

K.A. and B.T.G. conceptualized the study. K.A., H.R.W., and B.Z. designed and implemented the software tool. F.V., E.P., N.C., N.K., and B.T.G. performed data analysis. K.A. and B.T.G. prepared the final visualizations. B.T.G. and K.A. wrote the manuscript. All authors proofread the manuscript. B.T.G. and R.K. supervised the project.

