## Supplemental Tables 1-3 for "Hourglass, a tool to mine bioimaging data, uncovers sex-disparities in the IL-6-associated T cell response in pancreatic tumors"

**Table 1.** List of parameters obtained from stained PDAC tissue microarray core images using QuPath and CT-FIRE bioimage analysis softwares in combination with various postprocessing methods for image normalization

| **Parameter** | |
|---|---|
| H.Score | QuPath |
| Het.Score | Postprocessing |
| Num.1.Plus.Perc | QuPath |
| Num.2.Plus.Perc | QuPath |
| Num.3.Plus.Perc | QuPath |
| Num.Detections | QuPath |
| Num.Negative | QuPath |
| Num.Pos.per.um.2.Stroma | Postprocessing |
| Num.Pos.per.um.2.Total | QuPath |
| Num.Pos.per.um.2.Tumor | Postprocessing |
| Num.Positive | QuPath |
| Positive.Perc | QuPath |
| Pos.Pix.Perc.Stroma | Postprocessing |
| Pos.Pix.Perc.Total | QuPath |
| Pos.Pix.Perc.Tumor | Postprocessing |
| Pos.Pixel.Area.TU.Count | Postprocessing |
| Pos.Pixel.Area.STR.Count | Postprocessing |
| Tissue.Area | QuPath |
| SHG.NumFibers | CT-FIRE |
| SHG.Alignment | CT-FIRE |
| SHG.MeanLength | CT-FIRE |
| SHG.MeanStraight | CT-FIRE |
| SHG.MeanWidth | CT-FIRE |

**Table 2.** Terminology used in Hourglass workflow, definitions and examples.

| Term | Definition | Synonymous Terms / Examples |
|---|---|---|
| **Hourglass** | The pipeline described in this paper to process, analyze and explore multiparametric data. Also, the software application and corresponding R package. | |
| **Feature** | The variable of interest, a molecular marker. | e.g. Stain, Marker, Gene, Protein |
| **Parameter** | A readout/metric/measurement. Multiple parameters can be acquired for each feature. | e.g. Stained Area, Width, Number of Positive Pixels per mm2 |
| **Sample** | Each sample has a unique sample ID and used in **BySample (regional)** analysis.<br><br>Since there may be multiple samples per patient, this enables interrogation of differents levels of heterogeneity, and a direct comparison using a **paired** analysis. | also called:<br>Region of Interest (ROI),<br>Core (in tissue microarray (TMA)) |
| **Patient** | Individual that donated the sample.<br>Each patient is associated with unique case ID and 2-*n* samples, that may be averaged or aggregated for a **ByPatient (global)** analysis. | |
| **Annotation** | Any discrete label or description for feature+parameter and samples/patients. | |
| | **Sample/patient annotation:**<br>describes rows in multiparameteric data, used as main comparison, ie. clinical or histopathological data, transcriptomic classifications, etc. | e.g. Sex, SmokerStatus, Cancer Subtype, Stromal subtype, Overall Survival Time |
| | **Feature/parameter annotation:** describes columns in multiparameteric data (i.e. merged output from bioimage analysis). | e.g. Feature, Parameter, Is.Numeric, Keep.In.Analysis<br><br>e.g. for IL6_Number.of.Positive.Pixels, Feature = IL6 and Parameter = Number.of.Positive.Pixels |
| **Comparison** | Grouping samples and patients based on strata/labels in plots - can be a discrete or continuous variable. | e.g. compare sex differences for all features |
| | **Main comparison:** directly from user (leverages external data). See patient/sample annotations. | Note: continuous variables grouped into quantiles:<br>e.g. Age, 25-70 (low, intermediate, high) |

| | | |
|---|---|---|
| | **Custom comparison:** from multiparameteric data (leverages internal data), stratifies into discrete levels based on number of quantitiles indicated by user | e.g. Stain (IL6_Num.of.Pos.Pixels) becomes IL6_low, IL6_intermediate, and IL6_high |
| **Feature Set** | Grouping of features to compartmentalize and visualize subsets of data (usually biologically relevant ones). Users are required to select a parameter for each feature, standard and optional alternative. | e.g. Set name: Plasma cells; Set list: *IgG CD27, CD38, CD78, CD138, CD319*<br><br>can also group sets to make "super-sets", e.g. Set name: lymphocytes; Set list:Tcell,Bcell |

**Table 3.** Output of Hourglass: descriptions, example images, and relevant functions in R package.

| Output Name | Image (Example) | Description/use | R Functions |
|---|---|---|---|
| **Heterogeneity Barplot** | 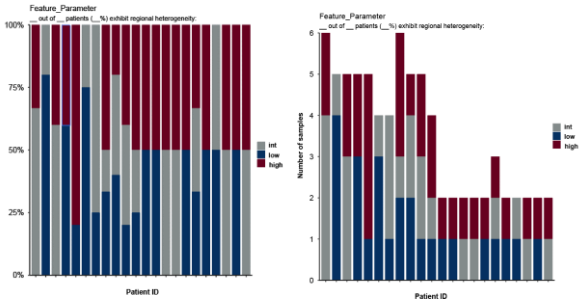 | Visualizes sample compositon (or expression levels within samples) in patients, indicative of intrapatient heterogeneity. | het_barplot.R: run_het_analysis, plot_het_barplot |
| **Survival Plot** | 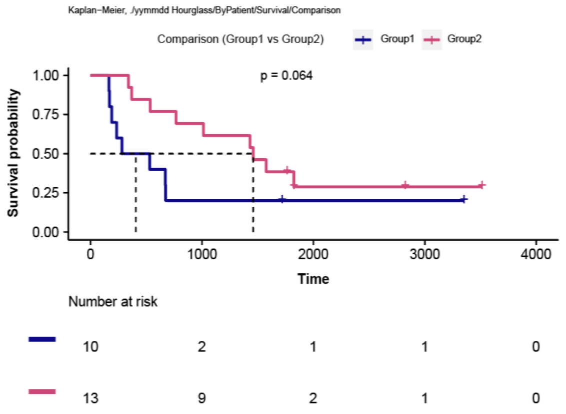 | Determines survival outcome via plotting of Kaplan Meier curve, logrank test, and tables containing patient counts. Only performed on patients (not samples). | survival_analysis.R: plot_surv_curve |
| **Paired Patient Slopegraph** | 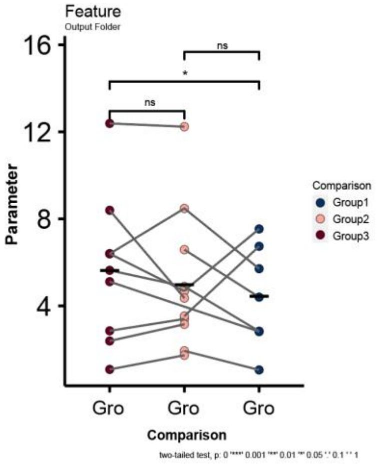 | Enables screening of any intra-patient differences that may be reflective of heterogeneity. Points are averaged values across strata per patient. Lines connect different averages within patients. | paired.R: get_paired_df, plot_indiv_paired |

<table>
<tr><td>Discrete Barplot</td><td>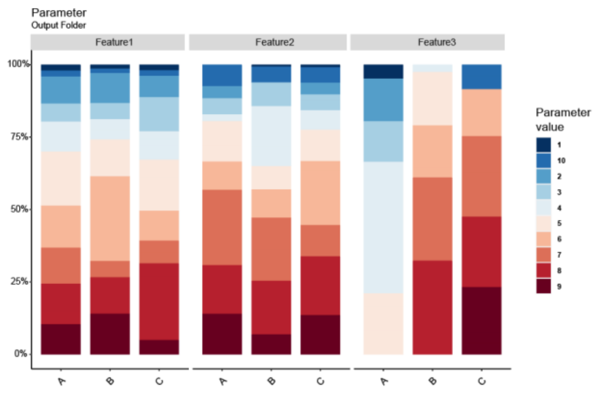
</td><td>Visualizes discrete values of a parameter in each stain, that is numeric in the input data but should be considered discrete. E.g. A parameter called "Heterogeneity Score" contains values 1-10 but should be "1", "2", .. "10".</td><td>discrete_barplot.R: plot_discrete_barplot</td></tr>
<tr><td>Overview Boxplot</td><td>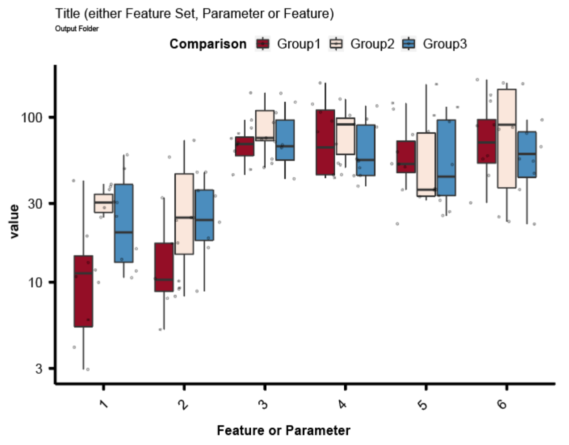
</td><td>Screen primary data for interesting relationships for all features in one plot.</td><td>boxplot.R: plot_overview_boxplot</td></tr>
<tr><td>Individual Boxplot</td><td>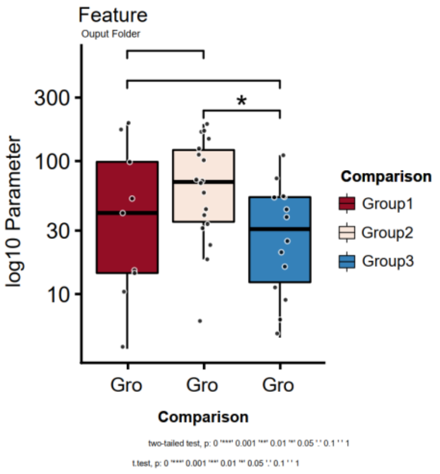
</td><td>Visualizes data points for each features/parameter combination and relevant statistics from comparing strata.</td><td>boxplot.R: plot_indiv_boxplot</td></tr>
</table>

| | | | |
|---|---|---|---|
| **Profile Barplot** | 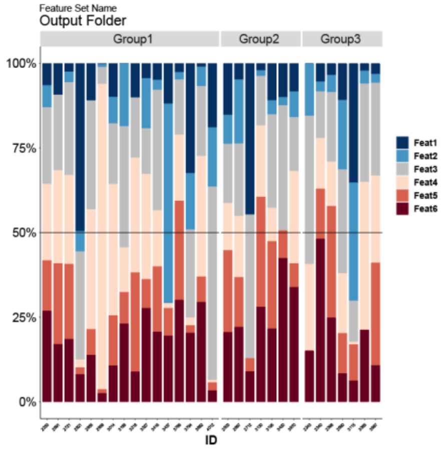 | Compare set of markers across different strata for each patient/sample. | profile_barplot.R: plot_profile_barplot |
| **Fold-change (FC) p-value Heatmap** | 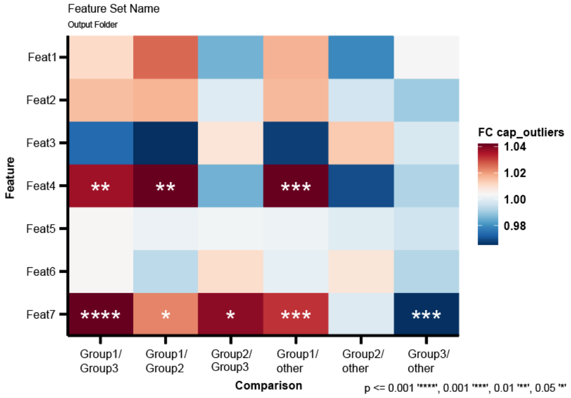 | Screen for patterns of fold-changes between strata in comparisons and overall significance seen in individuals boxplots. Fold-changes of strata are shown as color gradient and significance between two groups are presented as stars/numbers. | FC.pval.R: make_FC.pval_df, make_FC.pval_plot |
| **Expression Heatmap** | 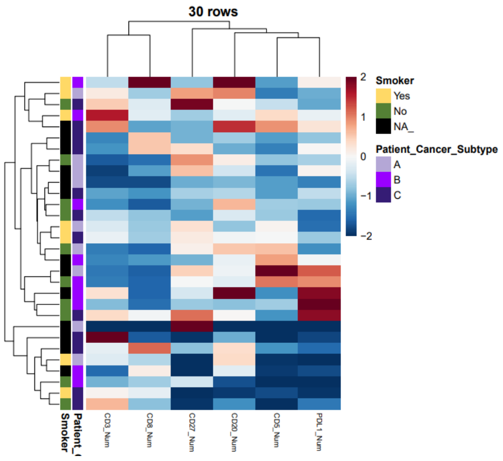 | Provides overview of patterns in stain data. When there are no NA/missing values (in imputed version for example), values will be clustered using unsupervised hierarchal clustering. | heatmap.R: plot_heatmap |

<table>
<tr>
<td>Correlation Plot</td>
<td>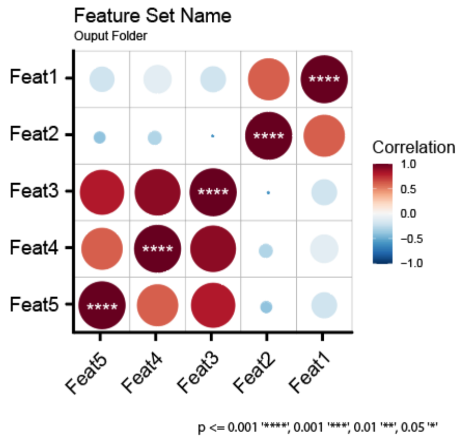
</td>
<td>Provides correlations between cell communities and their interactions, e.g. of an insight that can be drawn: correlations between fibroblast markers in one cancer subtype and anti-correlated with immune cells in another. Can only be performed on imputed (no missing values) dataset.</td>
<td>corrplot.R: plot_corrplotgg, plot_corrplot</td>
</tr>
<tr>
<td>Correlation Scatter Plot</td>
<td>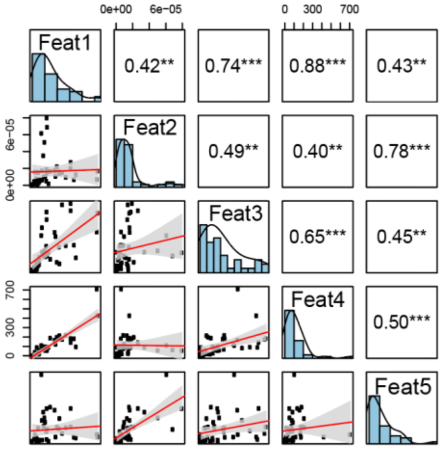
</td>
<td>Provides details of correlations, such as regression, scatter plots, correlation coefficients and significant and histograms of values used. Can only be performed on imputed (no missing values) dataset.</td>
<td>corrscat.R: plot_overview_corr_scatt</td>
</tr>
<tr>
<td>ID Table</td>
<td>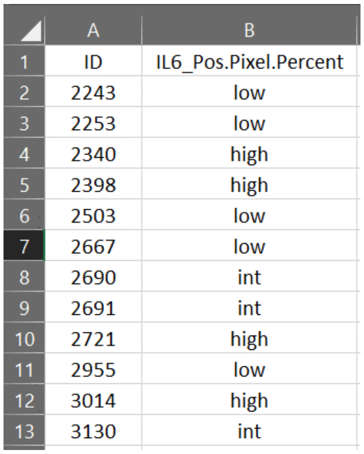

| | A | B |
|---|---|---|
| 1 | ID | IL6_Pos.Pixel.Percent |
| 2 | 2243 | low |
| 3 | 2253 | low |
| 4 | 2340 | high |
| 5 | 2398 | high |
| 6 | 2503 | low |
| 7 | 2667 | low |
| 8 | 2690 | int |
| 9 | 2691 | int |
| 10 | 2721 | high |
| 11 | 2955 | low |
| 12 | 3014 | high |
| 13 | 3130 | int |
</td>
<td>Provides sample/patient ID and strata used in comparison, useful for custom comparisons (where groups are assigned by Hourglass) and to see which samples/patients were included after filter inclusion/exclusion.</td>
<td></td>
</tr>
</table>

(1) Only Available for Feature Sets:
Profile Barplot, FC p-values Heatmap, Expression Heatmap, Correlation Plot, Correlation Scatter Plot
(2) Clustering/correlations can only be performed on data with no missing values (ie. imputed version, complete):
Expression Heatmap, Correlation Plot, Correlation Scatter Plot
