## Supplementary figures and images for "Hourglass, a tool to mine bioimaging data, uncovers sex-disparities in the IL-6-associated T cell response in pancreatic tumors"

### Supplemental Figures S1-S5

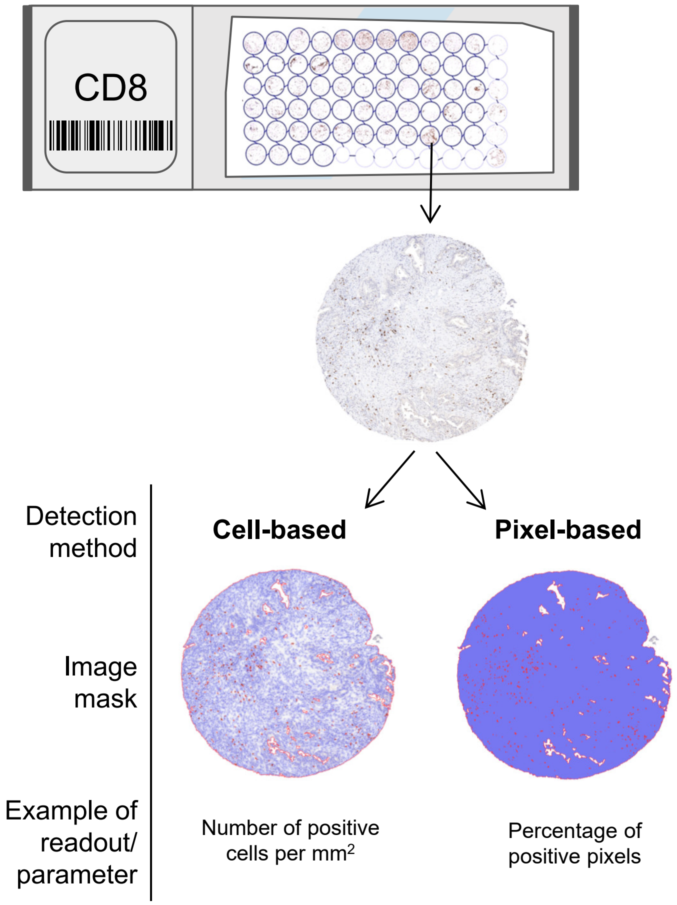

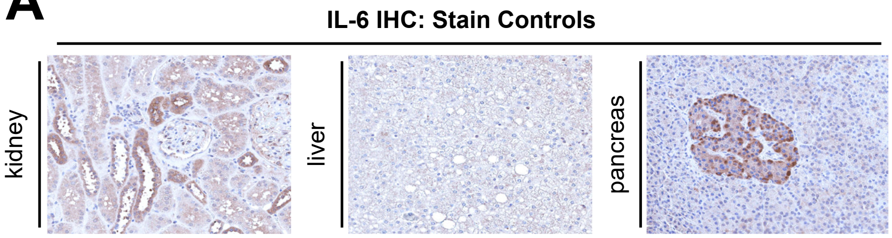

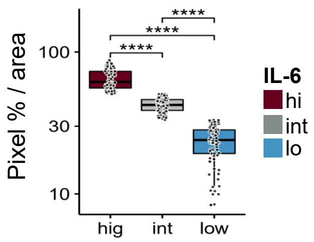

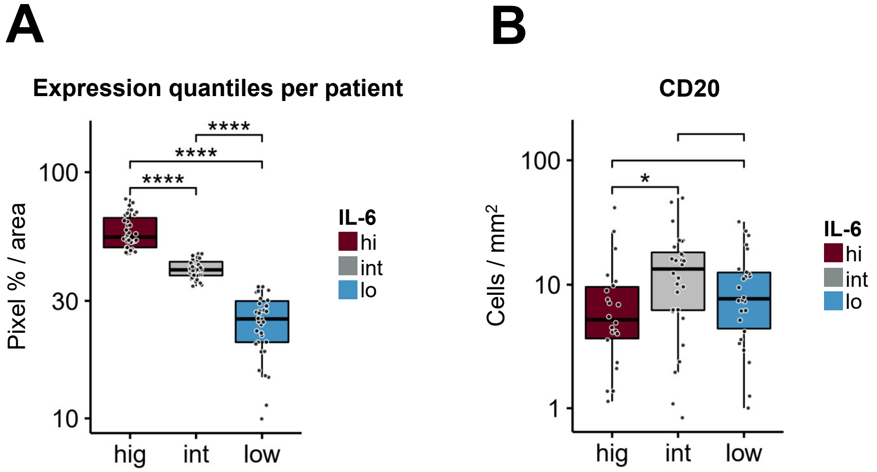

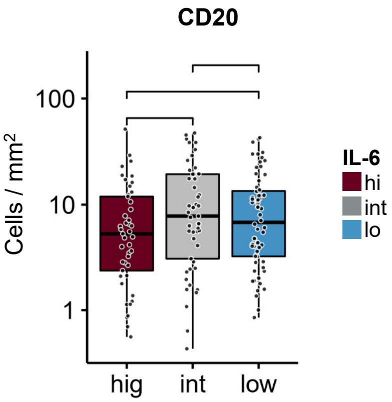


Figure S4

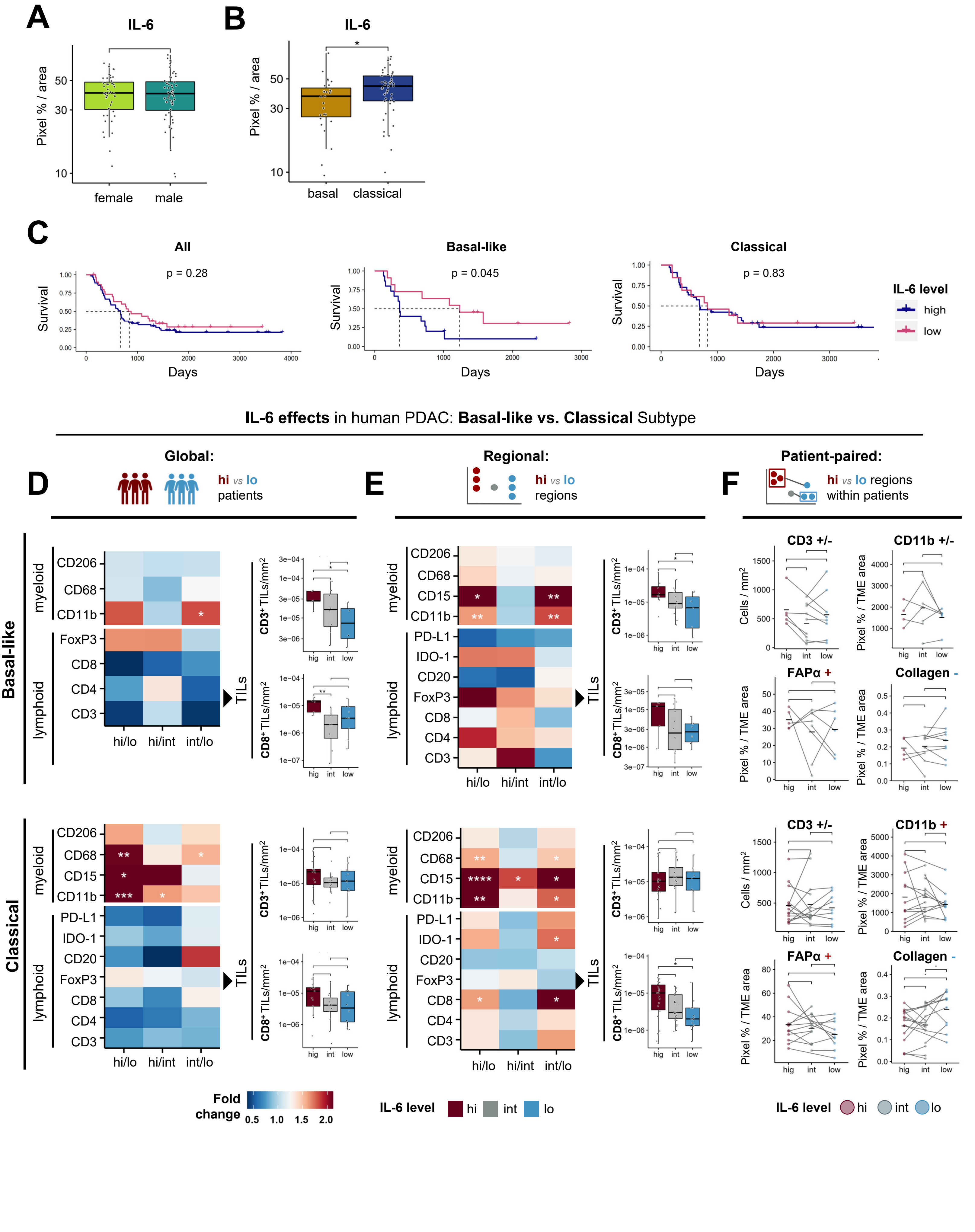
